# Dasatinib Plus Quercetin on Uterine Age-Related Dysfunction and Fibrosis in Mice

**DOI:** 10.1101/823229

**Authors:** Marcelo B. Cavalcante, Tatiana D. Saccon, Allancer D. C. Nunes, James L. Kirkland, Tamara Tchkonia, Augusto Schneider, Michal M. Masternak

## Abstract

Female reproductive function is negatively impacted by age. Animal and human studies show that fibrosis of the uterus contributes to gestational outcomes. Collagen deposition in the myometrial and endometrial layers is the main change related to uterine aging. Senolytic therapies are a potential option for reducing diseases and health complications related to aging. We investigated effects of aging and the senolytic drug combination of dasatinib plus quercetin (D+Q) on uterine fibrosis. A total of 40 mice, 20 young females (03-months) and 20 old females (18-months), were analyzed. Young (Y) and old (O) animals were divided into groups of 10 mice, with one treatment (T) group (YT and OT) and another control (C) group (YC and OC). Comparative analysis of *Pi3k/Akt1/mTor* and p53 gene expression among the 4 groups was performed to test effects of age and treatment on collagen deposition in uterine tissue. Uterine levels of microRNAs (miR34a, miR34b, miR34c, miR146a, miR449a, miR21a, miR126a, and miR181b) were evaluated. Aging promoted downregulation of genes of the *Pi3k/Akt1/mTor* signaling pathway (p=0.005, p=0.031, and p=0.028, respectively) as well as a reduction in expression of miR34c (p=0.029), miR126a (p=0.009), and miR181b (p=0.007). D+Q treatment increased p53 gene expression (p=0.041) and decreased levels of miR34a (p=0.016). Our results demonstrate a role for the *Pi3k/Akt1/mTor* signaling pathway in uterine aging and suggest for the first time a possible anti-fibrotic effect in the uterus of D+Q senolytic therapy.

## Introduction

The reproductive profile of women has been changing over the last few decades. Older maternal age by the first gestation and an increased number of pregnancies after 40 years of age are phenomena observed worldwide, impacting directly on gestational results [1]. Studies in humans and animals suggest a strong relationship between pregnancy loss and maternal age [2, 3]. In addition to advanced age, other gynecological conditions are related to fertility, such as polycystic ovary syndrome, leiomyomas, and endometriosis [4]. It is known that ovarian dysfunction is the major factor responsible for these poor reproductive outcomes, but other reproductive organs are involved in this complex process [2].

An increase in uterine volume with aging is common in some species of rodents, mostly due to endometrial cystic hyperplasia, as opposed to what occurs in menopausal women, in whom uterine atrophy is usually evident [3, 5]. The most obvious histological change in the aged uterus is the collagen deposition (fibrosis) in the muscle layers and stroma [3]. Mechanisms involved in this uterine fibrosis remain unclear [6]. Collagen deposition in tissues occurs as a result of chronic inflammatory processes involving several pathways: inflammatory interleukins, growth factors, caspases, oxidative stress products, and accumulation of senescent cells [6]. These chronic inflammatory pathways are also involved in undesired obstetric outcomes such as loss of pregnancies and preterm delivery [7].

The phosphoinositide 3-kinase (*Pi3k*)/ protein kinase B (*Akt*)/ mammalian target of rapamycin (*mTor*) pathway is an intracellular signaling mechanism that regulates several cellular functions, including cell growth, proliferation, differentiation, transformation, and survival, among others. Etiological processes underlying many gynecological conditions have not yet been completely identified. The *Pi3k/Akt1/mTor* and p53 signaling pathways appear to be involved in the pathophysiological mechanisms of gynecopathies including polycystic ovarian syndrome, premature ovarian failure, leiomyoma, endometriosis, and gynecological cancers [8-15]. This signaling pathway has also been implicated in fibrosis in different tissues, such as the kidney, lung, and liver [16-20].

*Pi3k* can be activated by binding of growth factors and steroid hormones to cell surface receptors, promoting conversion of phosphatidylinositol-4,5 bisphosphate (*PiP2*) to phosphatidylinositol-3,4,5 triphosphate (*PiP3*) [21]. The other components of the signaling pathway (*Akt-mTor*) are activated after *Pi3k*. *Pi3k* is a shared activator of two pro-fibrotic signaling pathways: *PAK2-Abl* and *Akt-mTor*. The activity of *Pi3k* is downregulated by enzymes phosphatases such as phosphatase and tensin homolog (*Pten*), which has been studied extensively with regard to mechanisms of gynecological cancers [11, 21]. The *Pi3k/Akt1/mTor* and p53 signaling pathways may also be jointly regulated by several microRNAs [17, 18, 22]. MicroRNAs are non-coding RNAs that act as transcriptional silencers and are involved in different cellular functions through post-transcriptional regulation of gene expression. Several microRNAs have been associated with the fibrosis process in different tissues (lung, liver, kidney, heart, skin) involving different mechanisms. Some of the most studied microRNAs in the process of fibrosis are: the miR34 family, miR126, miR181, miR21, miR146a, and miR 449 [22-24].

Targeting senescent cells with senolytic drugs might slow down or prevent fibrosis processes in different tissues and organs [16, 25-27]. Currently, quercetin (Q) and dasatinib (D), administered alone or in combination (D+Q), are the most studied senolytic drugs [28]. Different authors have reported anti-fibrotic effects of these drugs in tissues such as kidney, lung, and liver [25-27]. Quercetin is a flavonoid with antioxidant, anti-inflammatory, immunoprotective, and even anticarcinogenic effects [29]. Quercetin appears to have both estrogenic and antiestrogenic effects on the uterus, depending on the dose. However, studies about potential antifibrotic and senolytic effects of these drugs in the uterus are few, and there is no published study about effects of the D+Q combination on the uterus [30]. Dasatinib is an antineoplastic drug used to treat chronic myeloid leukemia and acute lymphoblastic leukemia [31]. Dasatinib’s anti-fibrotic effect has been ascribed during the last decade to its action on different signaling pathways such as *Pi3k/Akt1/mTor*, p53, and inflammatory pathways [25, 32, 33].

It is important to understand more completely the physiological mechanisms of uterine aging, as well as to discover therapies that delay this process. This could contribute to improvement in gestational outcomes. The aim of our study was to test the impact of aging on uterine fibrosis and the potential anti-fibrotic structural and molecular effects of the senolytic D+Q combination on uterine aging.

## Material and Methods

### Animal Studies

The Institutional Animal Care and Use Committee of the University of Central Florida approved all procedures in this study. BALB/c mice were obtained from NIA Office of Biological Resources and the NIA Aged Rodent Colonies and were maintained in a pathogen-free facility under temperature- and light-controlled conditions (22 + 2°C, 12h light/dark regimen) with free access to food and water. A total of 40 mice (females) were divided into four groups procedures: 1) Young controls: 3 month old mice given placebo treatment (YC; n=10); 2); Young treatment: 3 month old mice given dasatinib plus quercetin (D+Q) treatment (YT; n=10); 3) Old controls: 18 month old mice given placebo (OC; n=10); and 4) Old treatment: 18 month old mice given D+Q (OT; n=10).

The intervention (D+Q or placebo) was performed for 3 consecutive days every 2 weeks over a 10 week period. Dasatinib was purchased from LC Laboratories (Woburn, MA) and Quercetin from Sigma-Aldrich (St Louis, MO). Dasatinib (5 mg/kg) plus Quercetin (50 mg/kg) was prepared in a diluted solution composed of 10% ethanol (Sigma-Aldrich E7023; St Louis, MO), 30% polyethylene glycol 400 (Sigma-Aldrich 91893; St Louis, MO), and 60% Phosal 50 PG (Lipoid LLC, Newark, NJ). For both treatment groups (YT and OT), D+Q was administrated by oral gavage in 100–150µL and the control groups (YC and OC) received placebo solution by oral gavage.

### Histology/ Masson’s Trichrome Assay

Uterus samples were collected and dissected and small tissue fragments were placed into 10% neutral-buffered formalin immediately after necropsy and fixed for 24 hours. Thereafter, the samples were dehydrated in ethanol, clarified in xylol, and embedded in Paraplast. The samples were sectioned (5µm) and stained with a Masson’s Trichrome 2000™Stain kit (American Mastertech Scientific INC, McKinney, TX-USA) to detect deposition of interstitial collagen. The histological preparations were examined using a microscope (Axio Obeserver A1, Zeiss) with 4, 10, and 20x lenses.

### Western Blotting

The uterine tissue samples were homogenized in lysis buffer T-PER (Thermo Scientific, Waltham, MA, USA) containing a mixture of protease and phosphatase inhibitors. A total of 30µg protein was separated by electrophoresis and transferred to PVDF membranes. Nonspecific binding of antibodies was blocked with 5% milk in TBS-T for 1 hour at room temperature and probed with diluted antibodies specific for Collagen-1 (1:1000, ab88147, Abcam, Cambridge, UK) and *β*-actin (1:1000, G043, abm, Richmond, CA), followed by incubation with appropriate specific secondary antibodies. Immunoreactive bands were quantified by densitometry using the ImageJ software (Image Processing and Analysis in Java; U.S. National Institutes of Health Bethesda, MD, USA).

### RNA Extraction and Gene Expression

About 50 mg of the uterine tissue samples were homogenized with 1.0 mm zirconium oxide beads and 700 μL of Qiazol (Qiagen, Valencia, CA, USA). Total RNA was isolated using Qiagen RNeasy Mini Kit (Qiagen) columns following the manufacturer’s instructions. RNA concentration was measured by spectrophotometer and 1 μg of total RNA was converted into complementary DNA (cDNA) using an iScript reverse transcription kit (Bio-Rad Laboratories, Hercules, CA, USA). The cDNA samples were diluted to 10 ng/uL and stored at −20 C.

Real-time PCR using SYBR Green dye was used to evaluate uterine gene expression. The primers used in this study are listed in Table 1. PCR reactions were performed in duplicate, by adding 5μL of SYBR Green Master Mix (Applied Biosystems, Foster City, CA, USA), 0.4μL of forward and reverse primers (10μM solution), and 2μL of each cDNA sample, in a total volume of 20μL. Fluorescence was quantified using the Applied Biosystems QuantStudio™ 7 Flex System Fast RT-PCR system (Applied Biosystems™). For each assay, 40 PCR cycles were run (95 C for 3s and 62 C for 30s), and a dissociation curve was performed at the end of the reaction to verify the amplification of a single PCR product. Each assay included a negative control using RNase-free water.

**Table 1.**
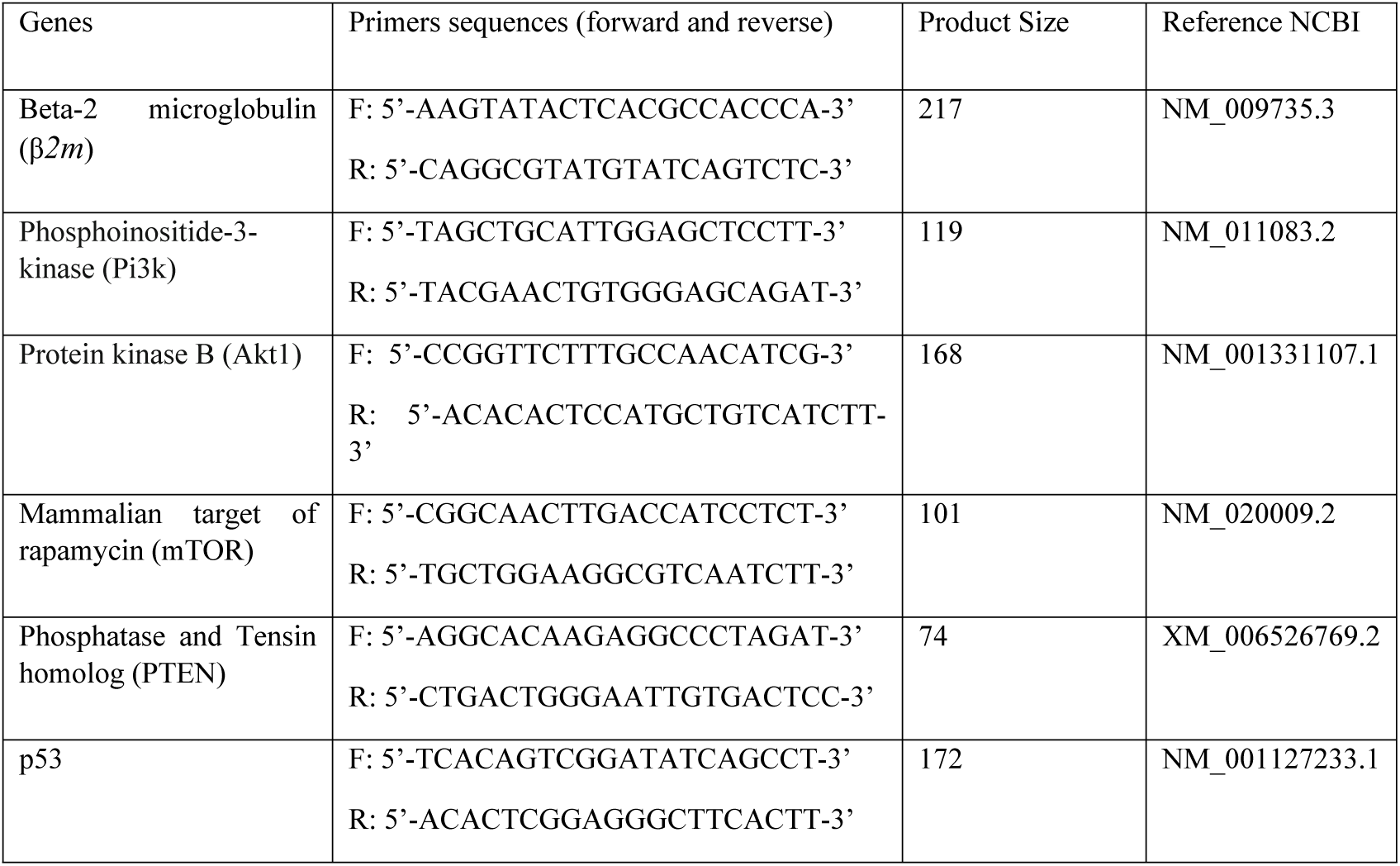
Primer pairs (forward and reverse) used in the experiment.

Data were normalized using beta-2 microglobulin (β*2m*) as a housekeeping gene. To calculate relative expression, the equation 2^*A*−*B*^/2^*C*−*D*^ was used, where *A* is the threshold cycle number of the first control sample of the gene of interest, *B* is the threshold cycle number in each gene of interest sample, *C* is the threshold cycle value of the first β*2m* in the control sample, and *D* is the threshold cycle number of β*2m* in each sample. This formula resulted in a relative expression of 1 for the first control sample, and then all the other samples were calculated in relation to the first sample. After that, the average of the YC group was calculated and used as a denominator for the other groups’ averages to calculate the fold change in gene expression compared to the control group [34].

### MicroRNA Expression

A total of 10ng of RNA was converted into complementary DNA (cDNA) using the TaqMan® Advanced miRNA Assays (Applied Biosystems™). The cDNA samples were diluted at 1:10 and stored at −20 C. The reactions were as *per* the manufacturer’s recommendation. Briefly, real-time PCR reactions were performed in duplicate, by adding 10 μL of TaqMan® Fast Advanced Master Mix (2X), 1 μL of TaqMan® Advanced miRNA Assay (20X), 4 μL of RNase-free water, and 5 μL of the diluted cDNA template to each reaction well in the plate. The total volume was 20 μL per reaction well. Fluorescence was quantified using the Applied Biosystems QuantStudio™ 7 Flex System Fast RT-PCR system (Applied Biosystems™). For each assay, 40 PCR cycles were run (95 C for 1s and 60 C for 20s). Each assay included a negative control using RNase free water. The TaqMan® Advanced miRNA Assays (Applied Biosystems™) used were: mmu-miR-16-5p (477860_mir), mmu-miR-146a-5p (478399_mir), mmu-miR-449a-5p (478561_mir), mmu-miR-21a-5p (477975_mir), mmu-miR-126a-5p (477888_mir), mmu-miR-34a-5p (478048_mir), hsa-miR-34b-5p (478050_mir), mmu-miR-34c-5p (478052_mir), and hsa-miR-181b-5p (478583_mir).

Data were normalized using miR16-5p as a housekeeping microRNA. To calculate relative expression, the equation 2^*A*−*B*^/2^*C*−*D*^ was used, where *A* is the threshold cycle number of the first control sample of the miRNA of interest, *B* is the threshold cycle number in each miRNA of interest sample, *C* is the threshold cycle value of the first miR16 in the control sample, and *D* is the threshold cycle number of miR16-5p in each sample. The formula resulted in a relative expression of 1 for the first control sample, and then all the other samples were calculated in relation to the first sample. After that, the average of the YC group was calculated and used as a denominator for the other groups’ averages to calculate the fold change in gene expression compared to the control group [34].

### Statistical Analysis

Statistical analysis was performed using GraphPad Prism 7 software (GraphPad Software Inc., La Jolla, CA, USA). Gene expression (mRNA), miRNA expression, and protein levels were compared between groups by 2-way ANOVA and p values for age, treatment, and its interaction are presented. When the interaction was significant, a multiple comparisons test was performed using Tukey’s test. Categorical variables were compared using the chi-square test. A p value lower than 0.05 was considered statistically significant.

## Results

During tissue dissection, it was noted that in old animals, 7 out of 20 mice (35%) had a dilated uterus. Among all old mice that had uterine dilation, 4/10 (40%) were from the D+Q group (OT) and 3/10 (30%) were from the control group (OC). There was no effect of treatment on the percentage of mice with a dilated uterus (p=0.639). Importantly, there were no cases of dilated uterus in young animals. The uterine tissue from mice with dilated uteruses was excluded from further experiments, which left remaining 6 old animals in the D+Q group (OT) and 7 old animals in the control group (OC).

Collagen deposition (fibrotic process) was observed in the muscular and endometrial uterine layers in histological analyses using Masson’s trichrome staining and confirmed by the presence of type 1 collagen in the uterine samples. Collagen deposition was significantly higher in old mice compared to young mice (age effect, p<0.001, Fig. 1) and there was no difference in fibrosis in treated groups compared to placebo (treatment effect, p=0.503, Fig. 1).

**Figure 1:**
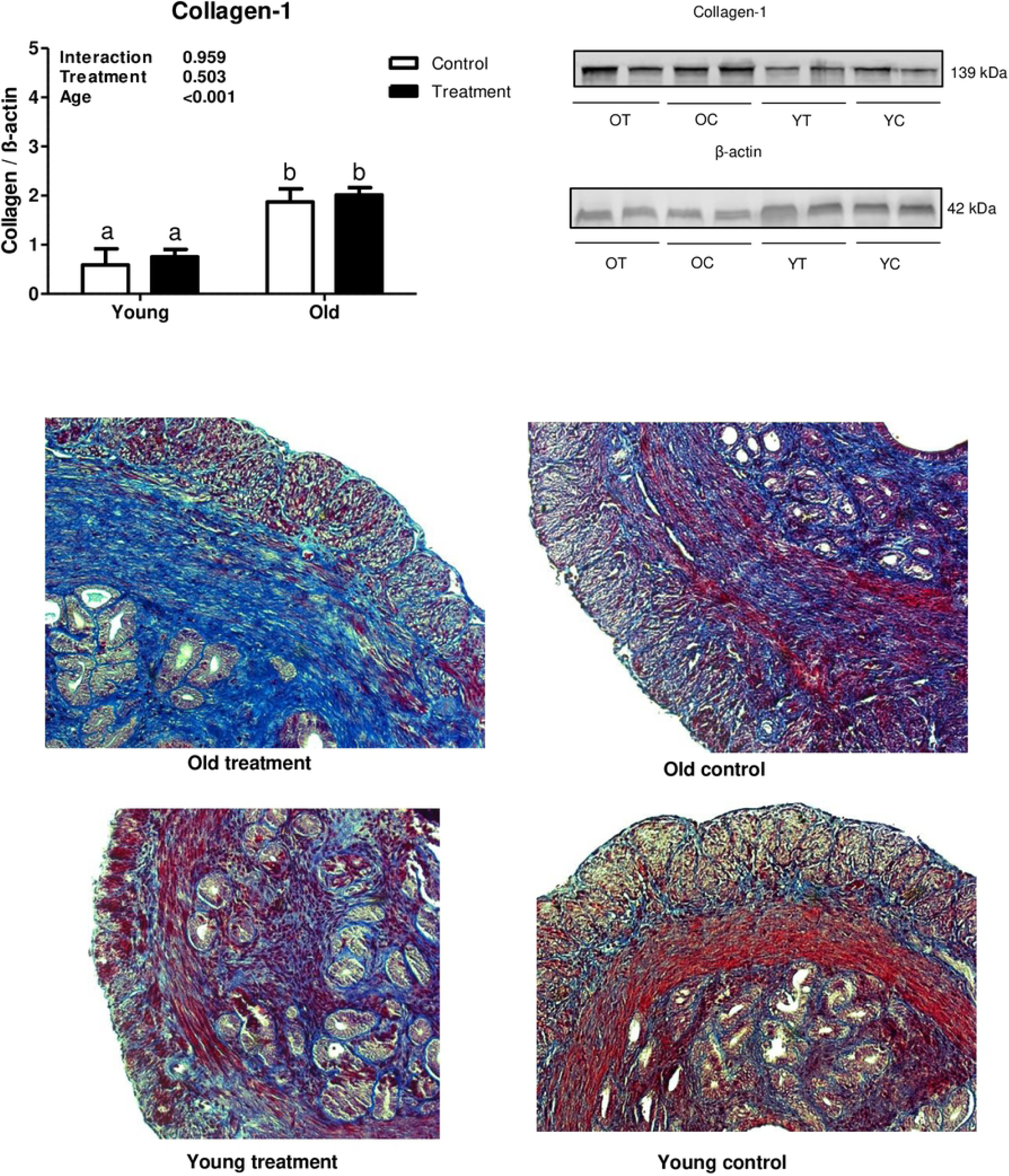
Uterine type 1 collagen deposition evaluation. (A) Collagen-1 statistical Western Blot analysis, letters indicate differences between groups (p<0.05), values were plotted as mean ± standard error of the mean. (B) Collagen-1 and β-actin Western Blot bands. (C) Representative Masson’s Trichrome stained images of uterine tissue. OT: old treatment; OC: old control; YT: young treatment; YC: young control.

Evaluation of uterine expression of different genes related to the *Pi3k/Akt1/mTor* signaling pathway revealed that aging was associated with inhibition of *Pi3k* and its downstream mediators, *Akt1* and *mTor*. The relative expression of *Pi3k, Akt1 and mTor* was significantly lower in old mice compared to young mice (p=0.005, p=0.031, p=0.028, respectively, Fig. 2A-C). However, there was no treatment effect on the expression of *Pi3k, Akt1, or mTor* (p=0.051, p=0.153, p=0.409, respectively, Fig. 2A-C). Regarding the gene expression of *Pten*, there was no effect of either treatment or age (p=0.394, p=0.064, respectively, Fig. 2D). Interestingly, p53 mRNA was upregulated with the D+Q treatment compared to control groups (p=0.041, Fig. 2E), while there was no aging effect (p=0.140, Fig. 2E).

**Figure 2:**
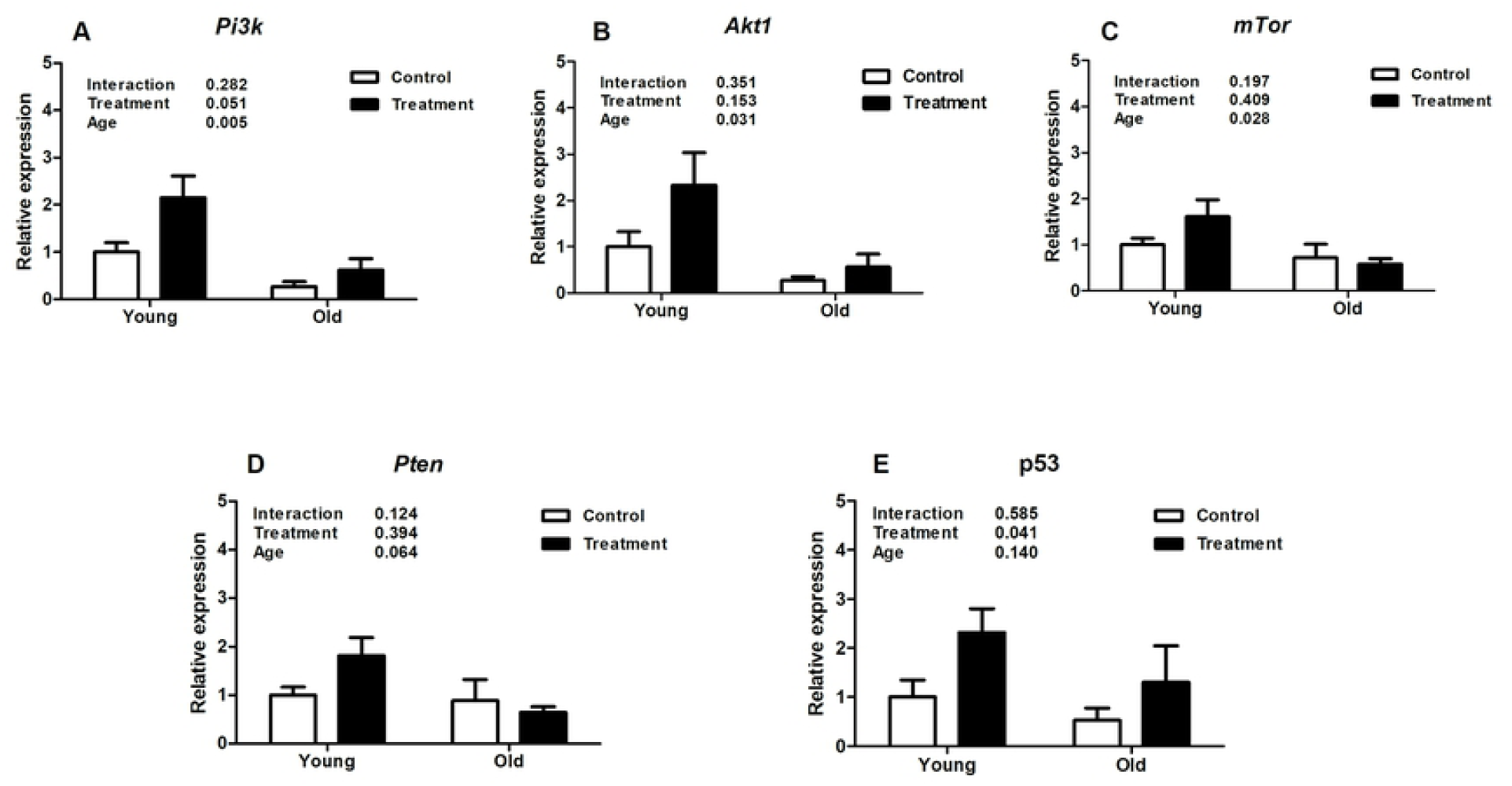
Analysis of relative uterine gene expression in treatment and control groups at different ages. A. Phosphoinositide 3-kinase (*Pi3k*). B. Protein kinase B (*Akt*). C. Mammalian target of rapamycin (*mTor*). D. Phosphatase and tensin homolog (*Pten*). E. p53. Values are shown as mean ± standard error of the mean. Two-way ANOVA was performed and the p values for age, treatment, and their interaction are presented (p<0.05).

Analysis of expression of different uterine microRNAs related to fibrosis pathways revealed that miR126a, miR34c, and miR181b were downregulated in old mice compared to young animals (p=0.009, p=0.029, p=0.007, respectively, Fig. 3G, C, H), however D+Q treatment did not affect expression levels in these miRNAs (p=0.958, p=0.352, p=0.446, respectively, Fig. 3G, C, H). Moreover, expression of miR34a was significantly decreased by D+Q treatment compared with the placebo control (p=0.016, Fig. 3A), while there was no uterine aging effect on miR34a expression (p=0.269, Fig. 3A). Additionally, aging and treatment did not affect the levels of miR146a (p=0.116 and p=0.067, treatment and age respectively, Fig. 3D), miR449a (p=0.632 and p=0.956, Fig. 3E), miR21a (p=0.416 and p=0.737, Fig. 3F), and miR34b (p=0.388 and p=0.490, Fig. 3B).

**Figure 3:**
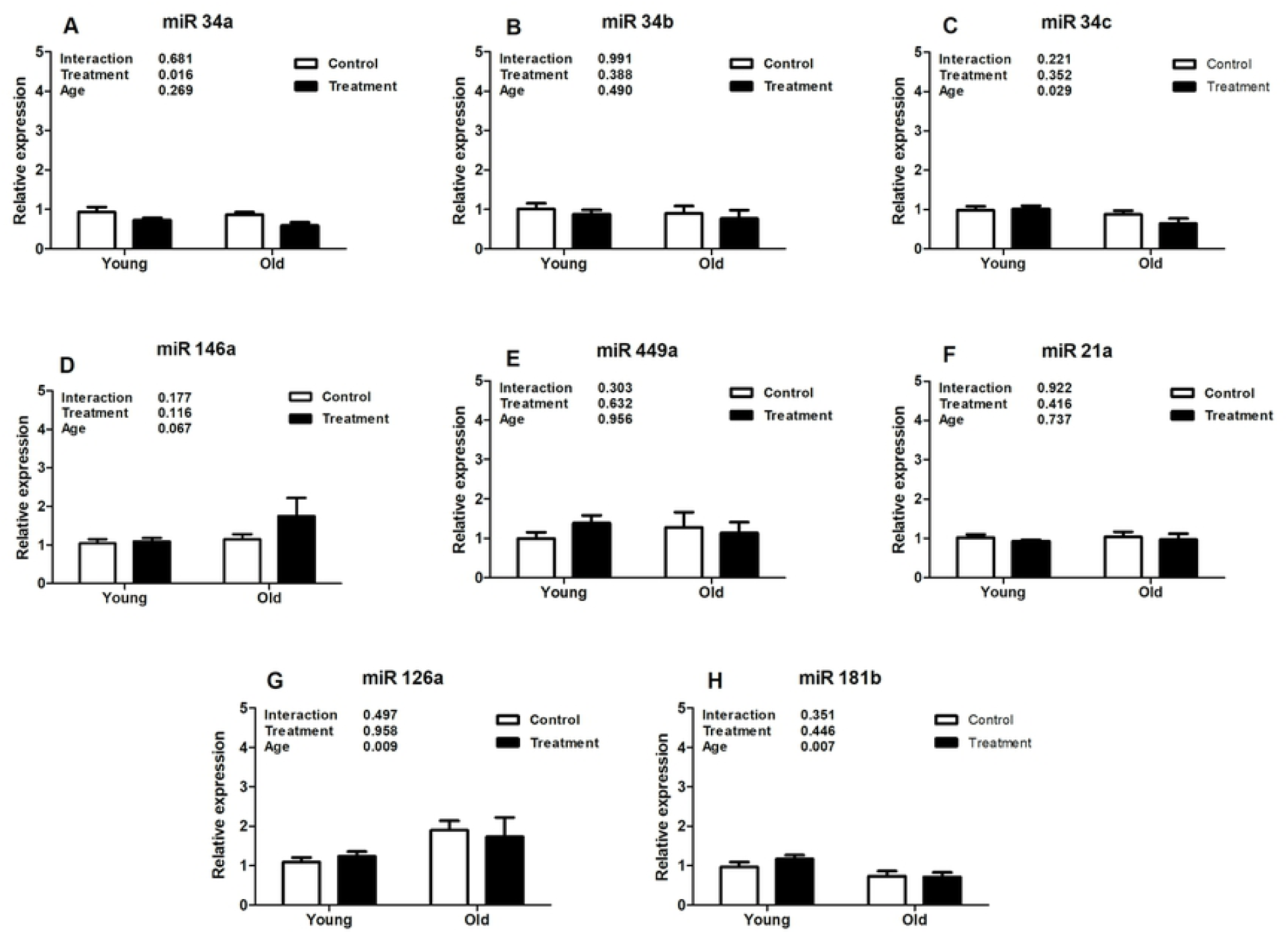
Analysis of relative uterine microRNA levels in the treatment and control groups at different ages. A. mmu-miR-34a-5p (miR34a). B. hsa-miR-34b-5p (miR34b). C. mmu-miR-34c-5p (miR34c). D. mmu-miR-146a-5p (miR146a). E. mmu-miR-449a-5p (miR449a). F. mmu-miR-21a-5p (miR21a). G. mmu-miR-126a-5p (miR126a). H. hsa-miR-181b-5p (miR181b). Values are shown as mean ± standard error of the mean. Two-way ANOVA was performed and p values for age, treatment, and their interaction are presented (p<0.05).

## Discussion

The main morphological changes observed during the mice uterine aging were increased uterine volume and fibrosis. In our study, dilated uterus was observed in 35% of the old mice, with no cases observed in any young mice. Interestingly, the D+Q treatment did not reduce the prevalence of uterine dilatation in old mice, while previously reported by Wilkinson *et al* a high prevalence of dilated uterus during aging (87%, 13/15 of old mice compared to 7%, 1/15 among young mice) was reduced by a high dose (42ppm) of rapamycin, a drug that inhibits the mTOR signaling pathway. However, in this study the authors continued the treatment for 13 months (from 9 to 22 months of age), which could prevent the development uterine dilatation rather than reverse it [5]. Despite that, D+Q treatment dose and time (10 weeks), which was started late in life in our study were not sufficient to observe a reduction in the prevalence of dilated uterus among the old mice. It might require long-term treatment starting in middle aged animals to observe possible preventive effects of D+Q on dilated uterus. Yet, due to upregulation of the *Pi3k/Akt/mTor* signaling pathway in endometrial hyperplasia and gynecological cancer, the samples with these pathological changes were removed from further genetic and histochemical analysis [9, 11].

The main feature of the uterine fibrosis process is collagen deposition, determined primarily by estrogen, a *Pi3k/Akt/mTor* signaling pathway activator [3, 35]. Therefore, uterine fibrosis observed in uterine aging is a chronic process, related to long and cyclical uterine exposure to estrogen. The *Pi3k/Akt/mTor* signaling pathway is downregulated in menopause due to a hypoestrogenic state [36]. Chong *et al* demonstrated that the change in gene expression in uterine muscle is dependent on exposure to female hormones, suggesting that the longer interval between menarche and first pregnancy worsens obstetric morbidity due to impaired myometrial function [37]. The literature reports an importance of *Pten* in improving longevity, due to its inhibitory action on *Pi3k* [38]. The increase in *Pten* gene expression could also contribute to *Pi3k/Akt/mTor* signaling pathway downregulation and consequently decrease collagen deposition [38].

Therefore, downregulation of the *Pi3k/Akt/mTor* signaling pathway at different points may be a useful treatment that can prevent the progression or even reverse fibrosis [39, 40]. One of the proposed therapies is single or combined use of D+Q that has an anti-fibrosis effect on different tissues such as lung, liver, kidney, and heart, but there is no report in the literature on the effect of these drugs on the uterine fibrosis process [25-27, 33, 41, 42]. Gao *et al* found that the cardiac anti-fibrotic effect of isorhamnetin, a quercetin methylated metabolite, occurs due to blockage of the *Pi3k/Akt/mTor* signaling pathway [43]. Cao *et al* found that quercetin is able to reduce the TGF-β-induced fibrotic process in human tubular epithelial HK-2 cells through miR-21 suppression and PTEN up-regulation [44]. Zhang *et al* in an *in vitro* study of the imatinib-resistant chronic myeloid leukemia cell line K562 (K562R^IMT^) demonstrated the inhibitory effect of dasatinib on the *Pi3k/Akt/mTor* signaling pathway and observed a slight upregulation of *Pten* at high doses of the drug [45]. Yilmaz *et al* observed that isolated use of dasatinib (8mg/kg/day for 21 days) was effective in reducing pulmonary fibrosis in an animal model by raising *Pten* levels [32]. Animal and human studies have shown that senolytic interventions provide a promising therapeutic possibility in cases of pulmonary fibrosis [28, 46, 47]. However, Roos *et al* reported that D+Q alleviated established vasomotor dysfunction in aged or atherosclerotic mice with no anti-fibrotic effect on the vascular intimal layer, which may suggest tissue and dose-dependent anti-fibrotic effects [48].

In our experiment, we showed the effect of aging on downregulation of the *Pi3k/Akt1/mtTor* signaling pathway in uterine tissue, but interestingly D+Q treatment did not promote an inhibitory effect on this pathway by either reducing *Pi3k/Akt1/mTor* gene expression or increasing *Pten* mRNA. As reviewed above, the effect of single or combined use of these drugs may be dose-, duration of treatment-, as well as tissue-dependent. In our study, the specific D+Q protocol used may explain the absence of a uterine anti-fibrotic effect due to the short duration of the intervention.

Importantly, our study indicated that D+Q treatment significantly increased the expression of p53 mRNA, the tumor suppressor gene that is related to a higher incidence of cancer in elderly people, which is not only due to a high frequency of mutant forms but its decline in its function with advancing age [13]. Several authors have described an increase in p53 levels with the isolated use of quercetin or dasatinib. Srivastava *et al* observed higher p53 expression in quercetin-treated tumor tissues [49]. The use of dasatinib also increased p53 acute myeloid leukemia stem cell gene expression [50]. p53 has also been described as a regulator of different microRNAs expression levels. The miR34 family includes the first miRNAs described as being regulated by p53 [51]. MiR34a has been found to be pro-fibrotic in various tissues such as lung, kidney, and heart [23, 52, 53]. The therapeutic inhibition of miR34a was effective in improving cardiac function after myocardial infarction in an animal model [54]. In our study, it was observed that the D+Q treatment promoted downregulation of miR34a, which could indicate a possible antifibrotic effect. Although the major regulatory pathway for miR34a expression is directly related to p53 levels, other p53-independent regulatory pathways are also known to be involved [55]. Therefore, p53-independent regulation of pro-fibrotic miR34a could contribute to the low expression of miR34a together with the high expression of p53 in our sample. Other members of the miR34 family were not impacted by D+Q treatment, and only miR34c was downregulated with aging. This is consistent with findings showing Pi3k pathway attenuation of the fibrotic process with age.

MiR21 is another fibrotic microRNA and it is regulated by *Akt* expression [22]. In our sample, although *Akt1* was downregulated with age, we did not observe similar downregulation of miR21a. Other pro-fibrotic microRNAs such as miR126a and miR181b were also downregulated in old uterine tissue, while miR146a and miR449a did not change in their expression either with age or treatment. The effect of age on serum and tissue microRNA levels has been tested in normal and long-lived (Ames dwarf) mice. Schneider *et al* observed an effect of age on levels of 22 microRNAs (out of 404 detected in sequencing) present in ovarian tissue from normal mice, and in 33 miRNAs from Ames dwarf mice [56]. Victoria *et al* also showed genotype-specific changes in the circulating levels of 21 miRNAs during aging. [57]. Therefore, this suggests that regulation of miRNA expression during aging is central to adaptation of body responses. As we have shown in our current studies, some miRNAs change with age in the uterus and others are regulated by D+Q treatment, further suggesting such a role.

## Conclusions

In summary, our findings suggest that uterine fibrosis is associated with the *Pi3k/Akt1/mTor* signaling pathway, with possible interaction and mediation of known pro-fibrotic microRNAs. Importantly, age-related fibrosis appears to be a slow and continuous process that might, over time, cause development of serious pathological complications, including those observed in our animals: a dilated uterus. Due to slow development of this age-related disease, D+Q senolytic therapy in the present protocol may not have been continued long enough for attenuating uterine collagen deposition. However, alteration of p53 mRNA and significant reduction of pro-fibrotic miR34a expression by D+Q suggest that implementing the intervention earlier in life and for a longer duration might provide protection from uterine age-related fibrotic changes. Conceivably, this might increase reproductive age as well as provide some protection against gynecological cancers. Based on these results, further, longer, and more mechanistic studies are required to determine whether *Pi3k/Akt1/mTor* pathway downregulation as well as inhibition of some microRNAs may provide new therapeutic targets to prevent uterine collagen deposition and, consequently, improve the reproductive performance of this organ in older females.

## Authors’ Contributions

MMM, MBC, TS, ADCN, AS, JLK, and TT contributed to experimental conception and design. MBC, TS, and ADCN performed the experiments. MBC, ADCN, and AS analyzed the data. MBC and ADCN wrote the first draft of the paper. All authors reviewed and approved the final manuscript.

